# Twitter and Mastodon presence of highly-cited scientists

**DOI:** 10.1101/2023.04.23.537950

**Authors:** Maximilian Siebert, Leonardo Maria Siena, John P.A. Ioannidis

**Affiliations:** Meta-Research Innovation Center at Stanford (METRICS), Stanford University, Stanford, CA, USA; Department of Public Health and Infectious Diseases, Sapienza University of Rome, Rome, Italy

**Keywords:** Twitter, Mastodon, social media, citations, scientific workforce

## Abstract

Social media platforms have an increasing influence in biomedical and other disciplines of science and public health. While Twitter has been a popular platform for scientific communication, changes in ownership have led some users to consider migrating to other platforms such as Mastodon. We aimed to investigate how many top-cited scientists are active on these social media platforms, the magnitude of the migration to Mastodon, and correlates of Twitter presence. A random sample of 900 authors was examined among those who are at the top-2% of impact based on a previously validated composite citation indicator using Scopus data. Searches for their personal Twitter accounts were performed in early December 2022, and re-evaluations were performed at 2 weeks, 4 weeks, and 2 months (February 6, 2023). 262/900 (29.1%) of highly-cited scholars had Twitter accounts, and only 9/800 (1%) had Mastodon accounts. Female gender, North American and Australia locations, younger publication age, and clinical medicine or social science expertise correlated with higher percentages of Twitter use. The vast majority of highly-cited author users of Twitter had few followers and tweets. Only 6 had more than 10,000 followers and none had more than 100,000. One limitation of our study is that it is possible that some accounts, especially with Mastodon, could not be detected. However, the study suggests that Twitter remains the preferred social media platform for highly-cited authors, and Mastodon has not yet challenged Twitter’s dominance. Moreover, most highly-cited scientists with Twitter accounts have limited presence in this medium.

## Introduction

Twitter has become a popular platform for researchers to communicate and share their work [1,2] and for influencers to comment on hot health and scientific matters. Currently Twitter has approximately 400 million users, of which half are estimated to be using it daily. Some users recently expressed dissatisfaction with Twitter after an ownership change and urged migration to another platform, Mastodon [3]. The latter saw an increase from 0.3 to 2.5 million users in the fall of 2022 and to 9 million in February 2023 [4,5].

While social media may shape scientific discourse and perceptions [2,6], it is unknown to what extent the most influential authors in the scientific literature are present in them and how intensively they use this medium. In 2017 only 1.12% of scholars with at least one publication since 2005 in the Web of Science had Twitter accounts; moreover, in that sample there was a highly skewed geographical location towards the United States of America (40%) [7]. However, the number of users increased by 20% during 2017-2022 and the COVID-19 pandemic gave impetus for more intensive use especially on biomedical and public health matters.

The current presence of publishing scientists who use Twitter would be interesting to know, especially for the scientists who are also the most influential in the scientific literature. Besides knowing what proportion of these scientists are present in Twitter and/or Mastodon, it would be useful also to know how active they are in these platforms, e.g. how many of them reach high levels of engagement. To answer these questions, we investigated the prevalence of top-cited scientists active on both social media platforms, the extent of migration to Mastodon, the intensity of social media activity of these scientists, and correlates of social media presence.

## Methods

The protocol for our analysis was pre-registered in OSF (https://osf.io/yc6z7/). We used the database of the top 2% most-cited scientists based on the citation impact of their work during 2021 [8]. The top-2% scientists in each of 174 scientific subfields are selected according to a previously validated composite citation indicator that incorporates information on citations, co-authorship and authorship positions – see references [9-11] for details on the construction and validation of the composite citation indicator. The composite indicator considers 6 metrics, including total citations; H index; co-authorship-adjusted Hm index; and numbers of citations to papers authored as single author; first or single author; and first, single or last author.

We limited our analysis to scientists who have published at least one paper in 2022, in order to exclude those who are dead or no longer active in the academic publishing field. We stratified top-cited authors according to their publication age and randomly selected 300 authors from those who first published before 1990, in 1990-2004, and after 2004 (n=900 total), so as to have good representation of relatively younger, mid-career, and older scientists.

The database of most-cited scientists that we used^7^ already included information on the scientific field, country, and publication age (year of first publication) for the sampled examined authors. Personal Twitter and Mastodon accounts for these 900 scientists such as gender and start dates of the accounts were manually independently searched online by two of the authors (LS & MS). Number of tweets and of followers were traced via twitter API v2 which allows researchers to access and interact with Twitter’s data and services, such as tweets, users, hashtags [12]. For Mastodon, the social media output and follower had to be collected manually.

Baseline searches happened in early December 2022, and re-evaluations were performed at 2 weeks, 4 weeks, and 2 months (February 6, 2023).

The primary outcomes were the proportion of researchers with Twitter and Mastodon accounts. Secondary analyses addressed changes compared to the baseline time point, and correlates of social media presence.

## Results

By February 6, 2023, 262/900 highly-cited scholars were on Twitter (29.1%; 95CI 26.2–32.1%). Over the 2-month time span, four researchers left Twitter and no one new joined. Nine highly-cited scholars (1%; 95CI 0.5–1.9%). were found on Mastodon. Seven joined in November, one in October and one in December 2022. All 9 still have active Twitter accounts.

The vast majority of active scholars were from North America or Europe. Female gender, North American and Australia locations, younger publication age, and clinical medicine or social science expertise correlated with higher percentages of Twitter use (**Table 1)**. Among the youngest scientists (those who started publishing in 2004 or later), more than a third (108/300, 36%) were present on Twitter, almost double the rate seen for older scientists (those first publishing before 1990). Clinical medicine was the most common discipline of work for Twitter users.

**Table 1.** Characteristics of highly-cited scientists on Twitter compared to those without an account by February 6, 2023. Percentages and 95% confidence intervals for categorical characteristics are given in brackets.

|  | With Twitter account<br>(n = 262) | Without Twitter account<br>(n = 638) | P-Value<br>(Fisher's exact test) |
| --- | --- | --- | --- |
| **Gender** |  |  |  |
| Male | 200 [76.3%; 70.7–81.3%] | 540 [84.6%; 81.6–87.4%] | .004 |
| **Region** |  |  |  |
| North America | 111 [42.4%; 36.3–48.6%] | 182 [28.5%; 25.1–32.2%] | <.001 |
| Europe | 97 [37%; 31.2–43.2%] | 216 [33.9%; 30.2–37.7%] | .41 |
| Asia | 25 [9.5%; 6.3–13.8%] | 202 [31.6%; 28–35.4%] | <.001 |
| Australia | 18 [6.9%; 4.1–10.6%] | 16 [2.5%; 1.4–4%] | .003 |
| Other | 11 [4.2%; 2.1–7.4%] | 22 [3.4%; 2.2–5.2%] | .72 |
| **First Year of Publication** |  |  |  |
| <1990 | 56 [21.4%; 16.6–26.8%] | 244 [38.2%; 34.5–42.1%] | <.001 |
| 1990-2004 | 98 [37.4%; 31.5–43.6%] | 202 [31.7%; 28–35.4%] | .11 |
| >2004 | 108 [41.2%; 35.2–47.4%] | 192 [30.1%; 26.6–33.8%] | .002 |
| **Field of Expertise** |  |  |  |
| Clinical Medicine | 104 [39.7%; 33.7–45.9%] | 157 [24.6%; 21.3–28.1%] | <.001 |
| ICT | 30 [11.5%; 7.9–15.9%] | 47 [7.4%; 5.5–9.7%] | 0.06 |
| Physics & Astronomy | 18 [6.9%; 4.1–10.6%] | 77 [12.1%; 9.6–14.9%] | 0.03 |
| Biology | 13 [4.9%; 2.7–8.3%] | 23 [3.6%; 2.3–5.4%] | 0.45 |
| Chemistry | 13 [4.9%; 2.7–8.3%] | 43 [6.7%; 4.9–9%] | 0.39 |
| Engineering | 12 [4.6%; 2.4–7.9%] | 72 [11.3%; 8.9–14%] | 0.003 |
| Biomedical Research | 12 [4.6%; 2.4–7.9%] | 33 [5.1%; 3.6–7.2%] | 0.84 |
| Social Sciences | 12 [4.6%; 2.4–7.9%] | 9 [1.4%; 0.6–2.7%] | 0.009 |
| EST | 12 [4.6%; 2.4–7.9%] | 77 [12.1%; 9.6–14.9%] | <.001 |
| Other areas | 36 [13.7%; 9.8–18.5%] | 100 [15.7%; 12.9–18.7%] | 0.53 |

ICT: Information and Communication Technologies; EST: Enabling and Strategic Technologies.

Most of the highly-cited author users of Twitter had few followers and tweets. Only 6/262 had more than 10,000 followers and none had more than 100,000. Only 9/262 had sent more than 10,000 Tweets. Number of tweets and number of followers were strongly correlated (**Figure 1**).

**Figure 1.**
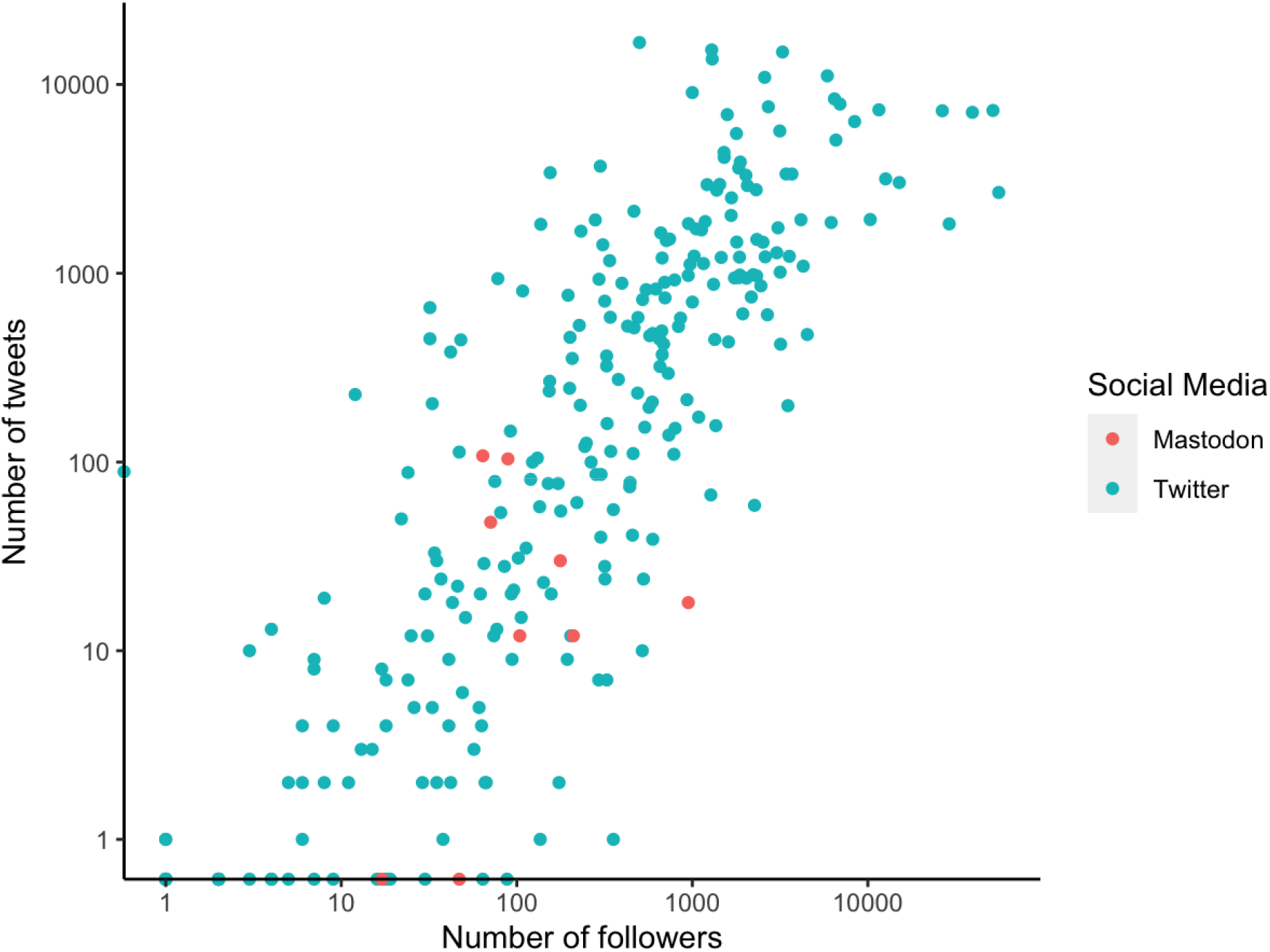
Number of followers and of posts by February 6, 2023 on Twitter and Mastodon. The points in blue represent Twitter users, in red Mastodon users.

## Discussion

Our results show that almost one in three highly-cited authors have a personal Twitter account and the rates are higher in younger scientists. This suggests that rates of Twitter presence for influential scientists are substantial. However, still the majority of influential scientists are not actively engaging with this social media tool. Moreover, even among those who do use Twitter, their presence is often limited in terms of the number of followers and tweets. Furthermore, despite a lot of discussion and occasional claims of leaving Twitter for Mastodon, migration to the latter remains negligible among highly-cited scientists, at least until February 2023. This lack of interest in Mastodon could be attributed to the platform’s difficulty to navigate between the different servers, having to start from scratch to gain an audience, and possibly other factors as well [13].

Our observed rates of Twitter presence are much higher than those documented in some earlier survey work covering all publishing scientists [7], but the selection criteria were different in the examined samples. Some assessments focusing on single disciplines have also shown high rates of Twitter engagement. For example, one study found in 2019 that Twitter utilization was 49.3% for Urology residency training programs in the USA and 34.1% of the physicians who were faculty in these programs were using Twitter [14]. We did not have sufficient power to explore whether specific subfields within larger fields (e.g. specific medical specialties) have greater engagement in Twitter than others. However, there is some evidence about differential engagement with Twitter across disciplines and this may include also the reasons this platform is used by different specialists [15]. Our examined sample had the advantage of covering all scientific fields, therefore giving an aggregate picture of the scientific workforce at different age groups.

Our approach has some limitations. First, finding researchers on Mastodon is more challenging compared to Twitter. Due to the platform’s open-source configuration comprising different servers and its search algorithm being less sensitive than the one of its opponent making it more difficult to find researchers on the platform. However, it is unlikely that any missed accounts would have changed the big picture: Twitter is still vastly more commonly used by scientists than Mastodon, similarly to the large difference that exists also in the relative use by the general public. Second, very young scientists, especially those still in training (e.g. doctoral and postdoctoral students, medical trainees, etc.), are not well represented in our sample, since it is extremely difficult for them to reach top-cited performance so quickly in their careers. It is conceivable that their participation in these social media may be even more prominent. Moreover, we did not attempt to separate different types of Twitter use, e.g. for expression opinions and debates versus more structured activities, such as education [16]. We also made no effort to judge the veracity, tone, sentiment and constructiveness (or lack thereof) of the tweets [17], since this is extremely difficult to do for people covering such a wide range of expertise. Even top-cited scientists may not necessarily use Twitter for scientific communication, but for more personal tastes and preferences. Fourth, we focused on personal accounts, but some scientists may also engage in Twitter through team or organization-level accounts. However, team and organization accounts do not express single individuals.

Overall, the study suggests Twitter presence is high among top-cited scientists, but most top-cited scientists still do not participate in this medium. Mastodon has not challenged Twitter’s dominance in the scientific community. Our study also highlights that powerful social media influencers with massive followings may have very little overlap with the group of hi6hly-cited authors in the scientific literature. Other authors have also drawn attention to the need to understand the background, expertise, and motivation of influencers who can shape scientific discourse [18]. There can be both rewards and risks in the engagement of academics in social media [19]. On one hand, social media engagement can reduce the feeling of isolation, facilitate interactions with a diverse range of researchers, and improve public engagement profiles; on the other hand, users can be exposed to personal abuse on these platforms. Social media presence is a distinct type of influence that is independent of and poorly correlated with other types of scientific impact, e.g. citations both for single papers [20] and for overall authors’ impact [21]. Therefore, it should be studied for its own merit and for the potential impact that it exerts.

